# Quantifying impacts of natural gas development on forest carbon

**DOI:** 10.1101/2025.06.23.661107

**Authors:** Elliot S. Shannon, Andrew O. Finley, Paul B. May

**Affiliations:** Department of Forestry, Michigan State University, East Lansing, MI, USA; Department of Statistics and Probability, Michigan State University, East Lansing, MI, USA; Department of Mathematics, South Dakota School of Mines and Technology, Rapid City, SD, USA

## Abstract

As energy demands continue to rise, energy production from sources including natural gas is expected to rapidly accelerate in the coming decades, potentially leading to substantial land-use changes. In the Appalachian region of the United States, natural gas development often occurs in forested areas, which can cause high levels of forest disturbance and loss. Here, we use nationwide forest inventory and remotely sensed data in a Bayesian model to quantify the impacts of natural gas development at fine spatial resolutions between 2008-2021. Based on well permit locations in the states of Ohio, Pennsylvania, and West Virginia, the analysis quantifies disturbance area, forest carbon loss, and opportunity cost with associated levels of uncertainty at the pixel-level. Overall, we estimate 10,854 ha of forest land were disturbed, resulting in 542,675 Mg (± 4,275) of forest carbon loss. The opportunity cost associated with these disturbances is estimated to be 575,246 Mg (± 30,774). Pixel-level estimates are generated for individual well sites, which can be aggregated to the county-level to highlight regional patterns. Specifically, we observe greater levels of disturbance in Northern West Virginia, while opportunity costs are greatest for large forested counties in Northeastern Pennsylvania. This study demonstrates the importance of quantifying balances and tradeoffs between energy production and forest ecosystem services, and provides important insights into the impacts of energy development on forest carbon.

## 1 Introduction

Natural gas energy production in the United States (US) has accelerated in recent years to meet rising energy demands. Associated land use changes, including those on forest lands, are an important consideration, as studies suggest that energy production will be the primary driver of land use change in the US through 2040 (Trainor et al., 2016; McDonald et al., 2009). At the same time, efforts to leverage carbon storage benefits of forests are also increasing (Favero et al., 2020). However, impacts of land use for natural gas infrastructure, including roads, pipelines, and well footprints, on forest ecosystems remain poorly quantified (Adams et al., 2011). In particular, the effects of natural gas development on forest carbon storage has received little attention. Therefore, assessing tradeoffs between natural gas development and forest carbon storage, including associated opportunity costs, is essential for informed decision-making.

The Appalachian states of Ohio, Pennsylvania, and West Virginia (Figure 4) are significant natural gas energy producers, with total production capacity expected to grow in the coming decades. Natural gas extraction from the Marcellus and Utica shale formations has a long history in the region (Slonecker et al.; Carter et al., 2011), reaching record production levels in 2021, when the region accounted for about one-third of the nation’s dry natural gas output (Grushecky et al., 2022b; U.S. Energy Information Administration, 2021). Advancements in horizontal drilling techniques have contributed to this significant increase, with further expansion anticipated in the coming decade (Jones et al., 2015). At the same time, much of this region is heavily forested, providing important environmental and economic benefits including forest carbon storage. Therefore, natural gas development in this region has potential to lead to substantial forest carbon loss (Slonecker et al.; Young et al., 2018).

While effects of land use changes associated with natural gas development have been previously studied, most work has investigated impacts on water, wildlife, and soils. One study examined land use change patterns and disturbance extents linked to over 4,000 oil and gas well pads in the Appalachian region of the US between 2007 and 2017 Grushecky et al. (2022a). During this period, natural gas production increased significantly, driving substantial land cover changes. The study found that disturbances from unconventional oil and gas development were greater than previously reported, with an average disturbance area of 5.6 ha per well pad, contributing to a total of 23,000 ha of affected land. Most land use changes were conversions of pre-existing forest land to grass-dominated systems.

A comprehensive review examined the impacts of natural gas production on biodiversity and ecosystem services Jones et al. (2015). The analysis highlighted key tradeoffs associated with energy development, including increased habitat fragmentation, noise and light pollution, the spread of invasive species, and water degradation. Another study quantified land use intensities of different energy sources, highlighting the substantial land use requirements for natural gas production Lovering et al. (2022).

Although the effects of energy development on forest structure, including canopy cover, have been studied (Adams et al., 2011; Young et al., 2018; Harris, 2020), few studies have quantified these impacts at regional scales. Notably, among 276 articles reviewed in (Jones et al., 2015), none specifically examined the extent to which energy development drives biomass loss from vegetation, including forests. However, an earlier analysis comparing the effects of wind, oil, and natural gas development on ecological indicators—including total biomass carbon stocks in Wyoming and Colorado—found that oil and natural gas production led to significantly greater overall carbon loss compared with wind energy (Jones and Pejchar, 2013). These effects were primarily due to the tendency for oil and gas wells to be sited in forested areas, unlike wind turbines.

Efforts to model status, trends, and change of forest carbon over space and time have received greater attention in recent years, often utilizing a combination of remotely sensed data and ground-based field measurements (Coops et al., 2025). Statistical methods leveraging sparse data to produce estimates for small areas, known as “small area estimation” (SAE), have also been developed. A series of recent studies using spatio-temporal SAE models identified areas of forest carbon density accumulation and loss for counties across the contiguous US (CONUS) from 2008–2021 (Shannon et al., 2024, 2025). Those studies found significant negative trends in forest carbon density in Appalachian region counties with potentially high natural gas development. These findings motivate the current study, which uses novel high spatial and temporal resolution models to better understand these trends.

We propose a Bayesian spatio-temporal SAE model to investigate impacts of natural gas development on forest carbon in the Appalachian region of the US between 2008-2021. The proposed model leverages nationwide forest inventory (NFI) data from the US Department of Agriculture Forest Service Forest Inventory and Analysis (FIA) program, as well as percent tree canopy cover (TCC) metrics from remotely sensed data to estimate forest carbon (defined here as carbon in aboveground live trees with diameter at breast height/root collar greater than 1 inch) at the locations of permitted natural gas wells. Estimates are produced at the pixel level, which are then aggregated to produce quantities of interest for wells and counties. The model proceeds via a two-stage framework, including a Bernoulli data model to predict probability of forest, and a latent Gaussian model to predict forest carbon conditional on the presence of forest. Threshold criteria are used to identify disturbed areas, where a covariate model is then developed to quantify opportunity costs. Opportunity cost captures both stored carbon lost through disturbance and new carbon the forest would have gained through growth if left undisturbed. Specifically, in areas of natural gas development, we seek to understand (i) what land area has been disturbed, (ii) how much carbon has been lost (iii) what is the opportunity cost in disturbed areas, and (iv) which forested areas have experienced the greatest carbon losses?

## 2 Results

The model detailed in Section 4.3 was fit using FIA and TCC data described in Section 4.2 to estimate pixel-level forest carbon for well aggregate locations outlined in Section 4.1 (Figure 4). Threshold criteria described in Section 4.4 were applied to identify disturbed pixels. Opportunity costs were estimated for disturbed pixel locations following methods outlined in Section 4.5. Additionally, to inform estimates of quantities of interest, including percent forest area disturbed, baseline estimates of forest area and carbon were generated from models fit using pixel-level TCC data for the entire study region.

Following disturbance threshold criteria described in Section 4.4, an estimated 10,854 ha of forest land were disturbed between 2009 and 2021, resulting in 542,675 Mg of forest carbon loss (534,296, 551,054 lower and upper 95% credible interval limits). This disturbed area represents approximately 0.07% of the total forested area within the states of Ohio, Pennsylvania, and West Virginia. The opportunity cost in these disturbed areas amounted to 575,246 Mg of carbon (514,928, 635,563).

Pixel-level carbon estimates were generated for all well aggregates, which are shown for a single well in Figure 1. National Agriculture Imagery Program (NAIP) imagery for disturbed well aggregates are regularly available over the time period, and highlight the progression of natural gas development. Annual estimates of total carbon for each well aggregate are compared for both disturbed and undisturbed scenarios, which provides insight into the opportunity cost associated with natural gas development.

**Figure 1:**
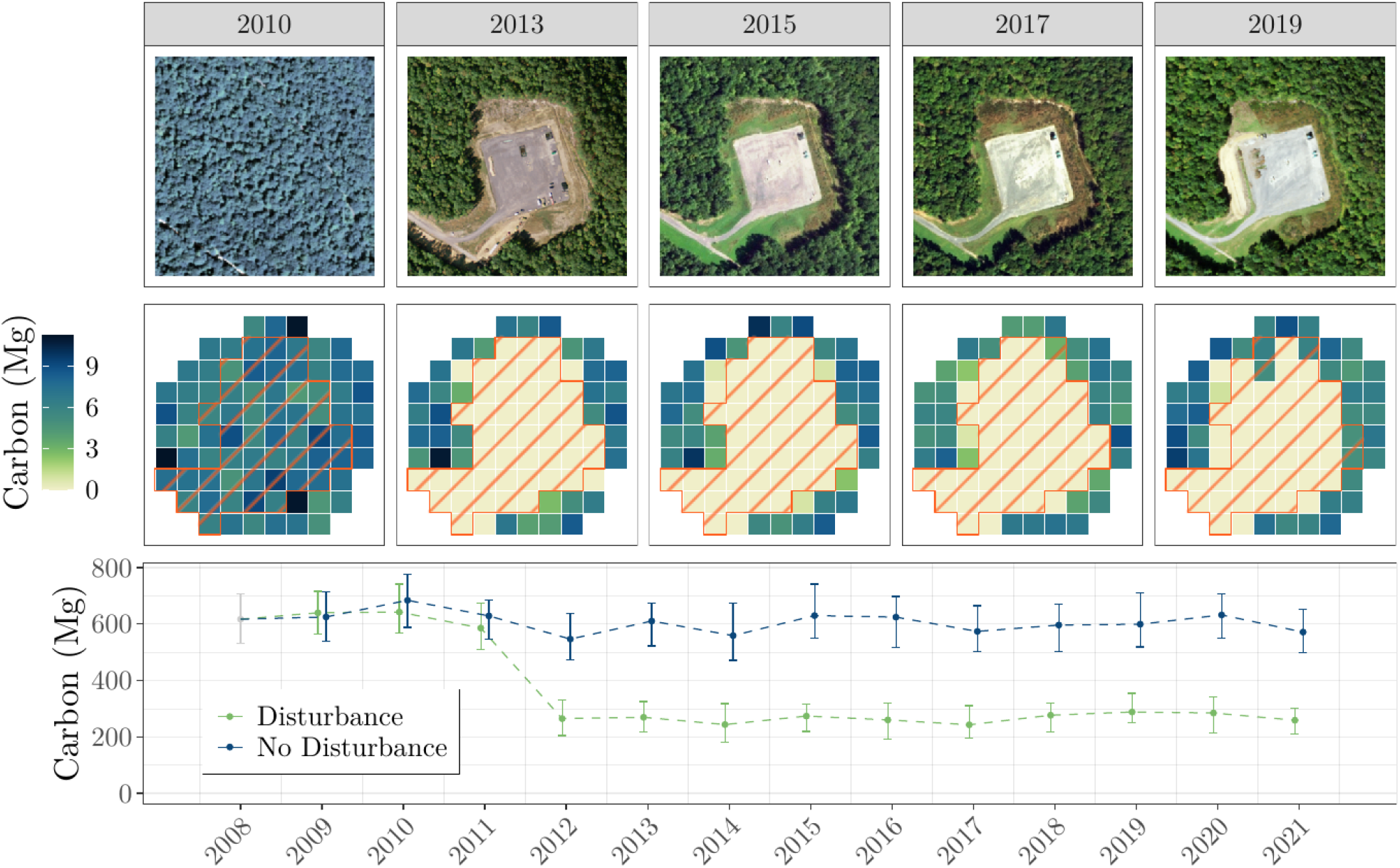
(top row) NAIP imagery displaying the time series progression of natural gas development for a well site in Tioga County, Pennsylvania. (center row) Posterior median pixel-level carbon estimates for years corresponding to available NAIP imagery. Disturbed pixels are denoted with an orange stripe pattern. (bottom row) Posterior median and 95% credible intervals of total carbon at this well site for both disturbed and undisturbed scenarios.

Pixel-level quantities and estimates are aggregated to the county-level and are displayed in Figure 2. Percent forest area is shown in Figure 2a, which highlights the distribution of forested areas in the region. Percent of total county area comprised of well aggregates is displayed in Figure 2b, which visualizes the distribution of natural gas development in the region. Insights from Figures 2a and 2b support patterns of percent forest area disturbed shown in Figure 2c, where heavily forested counties associated with significant natural gas development experience the highest levels of forest disturbance. Doddridge County, West Virginia, exhibited the greatest disturbance levels, amounting to approximately 0.7% (521 ha) of the total forest land in the county. Opportunity costs associated with disturbed locations are aggregated to the county-level and displayed in Figure 2d. Here, total opportunity costs are highest for the large, heavily forested counties in Northeastern Pennsylvania, with Susquehanna County, Pennsylvania, exhibiting the greatest total opportunity cost of 49,255 Mg of carbon (46,193, 52,317).

**Figure 2:**
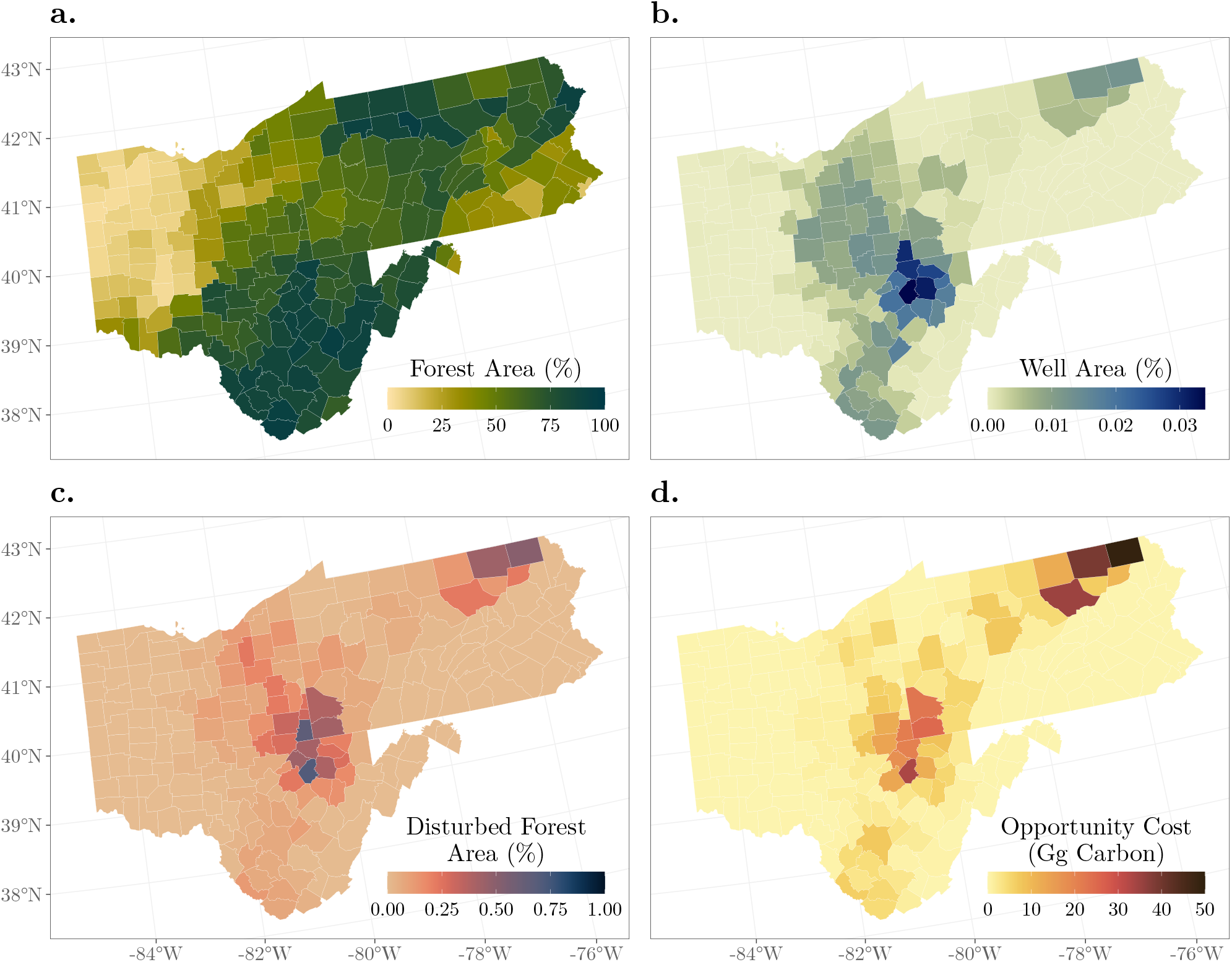
(a) Posterior median estimates of percent forest area for counties in Ohio, Pennsylvania, and West Virginia. (b) County-level percent well area, representing the percent of the total county area occupied by well aggregate areas. (c) Posterior median estimates of percent forest area disturbed at the county-level. (d) Posterior median total opportunity costs estimated at the county-level.

Baseline pixel-level carbon estimates for Susquehanna County, Pennsylvania, along with opportunity costs at disturbed locations are displayed in Figure 3. Estimates shown in Figure 3a highlight the complex distribution of forest in the county, and the focal area shown in Figure 3b provides insight into the density and impact of natural gas development in the area through pixel-level opportunity cost estimates.

**Figure 3:**
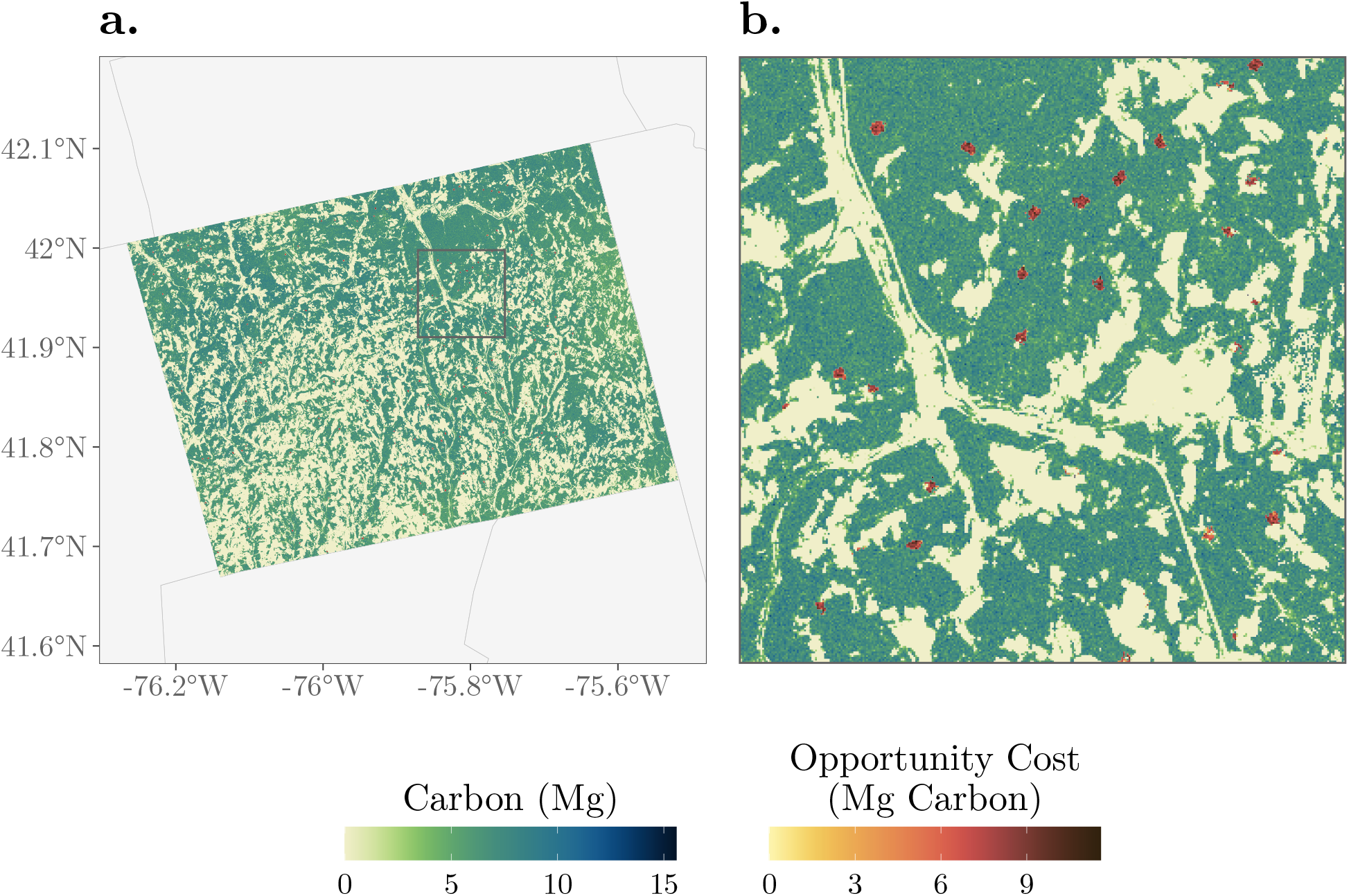
(a) Baseline pixel-level posterior median forest carbon estimates for Susquehanna County, Pennsylvania, with posterior median opportunity costs mapped for disturbed pixel locations. (b) Focal area highlighting the fine distribution patterns of forest carbon in the county, as well as the density and magnitude of opportunity costs at disturbed pixel locations.

**Figure 4:**
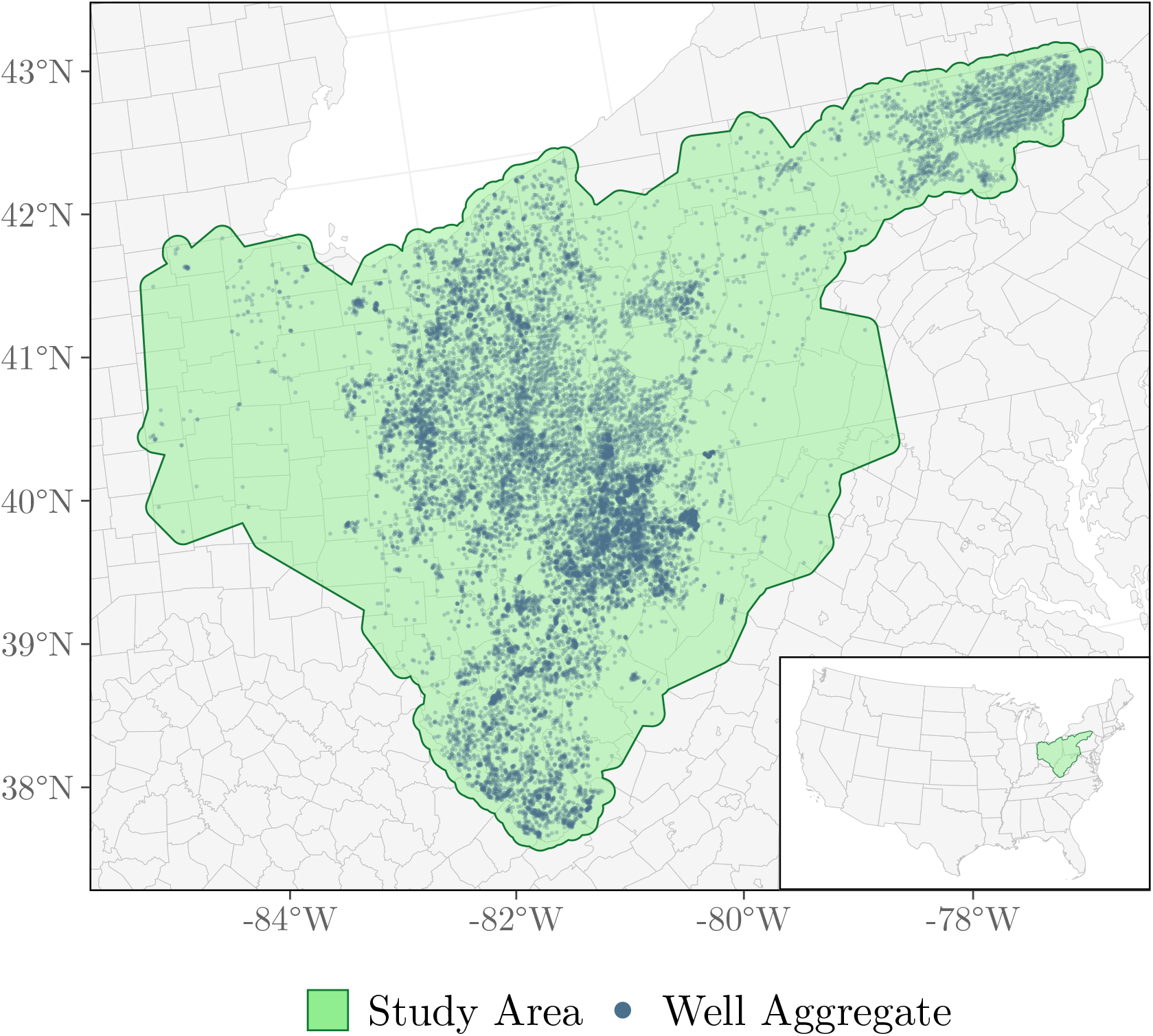
Study area generated as a 10 km buffer surrounding the convex hull of aggregated well permit locations. An inset map displays the study area location within the CONUS.

## 3 Discussion

Using NFI and remote sensing data, we developed a Bayesian spatio-temporal SAE model to quantify the impacts of natural gas development on forest carbon. Pixel-level estimates of forest carbon were generated for permitted natural gas well locations between 2008-2021 in the US states of Ohio, Pennsylvania, and West Virginia, and quantities of interest including disturbed forest areas and opportunity costs were derived. Estimates were also aggregated from the pixel-level to the county-level to identify regional patterns and trends.

Overall, an estimated 10,854 ha of forested lands were disturbed between 2009-2021.

This value is less than that reported by Grushecky et al. (2022a), who identified 23,000 ha of land affected by oil and natural gas development in the region between 2007-2017. However, their study considered all land-cover classification changes within buffered well pad areas, and did not directly estimate the probability of forest presence or disturbance during the study period. By considering probability of forest estimated at the pixel-level using NFI and remotely sensed TCC data, our estimate reflects pixel-level changes in forest structure rather than land-cover classifications. However, as outlined in Section 4.4, our estimate of total forest area disturbed is sensitive to the selected threshold value for probability of disturbance over the study period. Here, we aim to balance identification of marginal disturbances associated with natural gas development while filtering minor disturbances associated with natural forest processes. In this way, our criteria for identifying disturbances may differ from other approaches such as those based on land-cover classification or imagery. At the county-level, areas including Eastern Ohio, Northern West Virginia, and Northeastern Pennsylvania, exhibited the highest rates of disturbance on forested lands. These patterns reflect both the effects of increased energy development in heavily forested regions (Northern West Virginia) and lesser energy development in more moderately forested regions (Eastern Ohio).

Within disturbed well permit locations, an estimated 542,675 Mg of forest carbon was removed, contributing to an opportunity cost of approximately 575,246 Mg of carbon. This opportunity cost reflects both carbon that is directly removed at disturbed locations, as well as predicted carbon which would have otherwise accumulated had disturbance not occurred. This difference is highlighted for a single well in Figure 1, where the difference in total carbon estimates in 2021 for the ‘Disturbance’ and ‘No Disturbance’ scenarios represents the opportunity cost for this well. For locations that are undisturbed, the opportunity cost is zero. Similarly, locations experiencing significant disturbance (such as in clearcut areas that had been previously heavily forested), the opportunity cost is high. The autoregressive moving average (ARMA) model described in Section 4.5 provides only a simple estimate of relatively constant forest structure, and future work could incorporate more complex models of forest growth to better estimate carbon in undisturbed scenarios.

While the current study demonstrates a novel and sophisticated statistical model to investigate impacts of energy development on forests, a number of limitations persist. First, the primary data considered here include only permitted natural gas locations rather than verified areas of energy development. In many cases, these well permits did not result in clear well construction, as validated by NAIP imagery, which could lead to mis-attribution of forest carbon loss. Hence, we emphasize the results here represent only disturbances and opportunity costs associated with permitted well aggregate locations, and may not reflect actual impacts from well development alone. Second, criteria for determining the occurrence of disturbance for specific locations as described in Section 4.4 does not explicitly identify the timing of single unique disturbances. Therefore, we are unable to describe the temporal trends of natural gas development in the region. Future work could leverage methods in time series and change-point detection to associate disturbance events with specific time periods.

The impacts of natural gas development in forested regions are substantial, and methods to quantify their magnitude and identify spatial distributions and patterns are vital for meeting rising energy demands while maintaining forest ecosystem services. In the context of climate change, the carbon storage benefits of forests provide an opportunity to sequester greenhouse gases, while energy sources such as natural gas come with their own climate effects. Navigating this balance will require new methods and the incorporation of data from a variety of sources. The integration of these data and development of modeling techniques requires careful consideration to uncertainty quantification and methods for scaling estimates to varying levels of interest. Here, we demonstrate a modeling framework that can accommodate a variety of data sources at different scales to provide insights for future decision-making in energy development and forest management.

## 4 Methods

### 4.1 Study Area and Well Data

The US states of Ohio, Pennsylvania, and West Virginia are located in the Appalachia region of the Northeast US, where significant natural gas well development has occurred over the Marcellus and Utica shale formations in recent years. Here, publicly available permitted natural gas well locations between the years of 2008-2021 were obtained from the Ohio Department of Natural Resources, the Pennsylvania Department of Environmental Protection, and the West Virginia Department of Environmental Protection. Following estimated disturbance areas reported in Grushecky et al. (2022a) associated with natural gas well pads, circular buffers were generated around each well, with areas of 4.7, 6.2, and 4.4 ha for well permit locations in Ohio, Pennsylvania, and West Virginia, respectively. Overlapping buffer regions were then merged, resulting in 19,111 aggregated well permit areas ranging in size from 4.4 to 815.6 ha in size. The total study area was then taken to be a 10 km buffer surrounding the convex hull of these aggregated well permit areas (referred to as ‘well aggregates’ in the main text) (Figure 4).

### 4.2 Forest Data

The FIA program is tasked with the design and collection of NFI data across the US, whereby a network of approximately 300,000 inventory plots are visited on a rotating basis every 5-10 years (Bechtold and Patterson, 2005; Westfall et al., 2022). Forest variables of interest, including derived quantities such as live forest carbon density (Mg/ha) (simply referred to as ‘forest carbon’ in the main text) are measured at inventory plots, and serve as primary data for subsequent SAE models. Additionally, FIA determines whether each inventory plot measurement occurs on forested or non-forestland, and this information can be used to build Bernoulli models of forest presence or absence. In this study, 99,345 FIA plot measurements within the study area (Figure 4) were used to predict the presence of forest in subsequent Bernoulli models between 2008-2021, while a subset of 56,881 forested FIA plot measurements were used to estimate forest carbon conditional on forest presence.

Remotely sensed percent tree canopy cover (TCC) is produced by the USDA Forest Service as part of the National Land Cover Database (NLCD), which represents the percentage of land area covered by forest canopy (Housman et al., 2023). TCC is a remotely sensed data product derived from multispectral Landsat observations, and is available as annual raster images with a 30 m pixel resolution across the CONUS between 2008-2021. Here, pixel-level TCC values coincident with FIA plot measurements and well aggreagtes were used as covariates in subsequent models to inform estimates of forest presence and forest carbon. Additionally, TCC values for the entire study region were used to predict baseline estimates of forest area and forest carbon from fitted models.

### 4.3 Model

We used the latent Gaussian model (LGM) proposed by May and Finley (2025) to estimate pixel-level forest carbon within well aggregates. Let *y*(**s**, *t*) ≥ 0 be the forest carbon at location **s** at time *t*. To accommodate the significant probability mass occurring at zero in the forest carbon data model, a binary process *p*(**s**, *t*) is defined where *p*(**s**, *t*) = 1 ⇔ *y*(**s**, *t*) > 0. The LGM is then

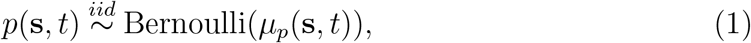

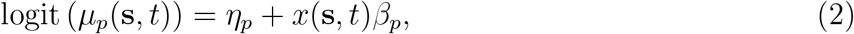

where *η*_*p*_ is an intercept term and *β*_*p*_ is a regression coefficient corresponding to TCC, denoted *x*(**s**, *t*). The forest carbon *y*(**s**, *t*) is then modeled as

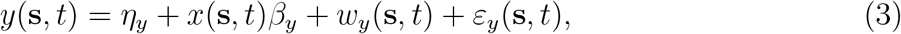

where 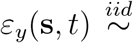 Normal(0, *τ* ^2^) is a normally distributed random error term, *η*_*y*_ is an intercept term, *β*_*y*_ is a regression coefficient corresponding to *x*(**s**, *t*), and *w*_*y*_(**s**, *t*) is a spatio-temporal random intercept following a mean zero Gaussian process. A final equation for forest carbon is then written as *y*(**s**, *t*) *× p*(**s**, *t*), which we denote as *z*(**s**, *t*).

A Gaussian process specification for *w*_*y*_(**s**, *t*) is represented as a sum of independent component Gaussian processes as in May and Finley (2025). These components allow for process variation to be represented at different scales. Here, we specify two components for the process, allowing for both short and long range dependencies in the spatial and temporal domain. Specifically, for a generic mean zero spatio-temporal Gaussian process *w*(**s**, *t*), we write *w*(**s**, *t*) ~ *GP* (0, *C*(·, ·)), where *C*(*δ*_*s*_, *δ*_*t*_) is a covariance function depending on the spatial and temporal lags *δ*_*s*_ and *δ*_*t*_, respectively. Implementing the dual component covariance structure as proposed in May and Finley (2025), we have

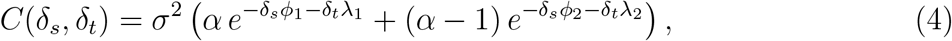

where *ϕ*_1_ > 0 and *λ*_1_ > 0 are the short range spatial and temporal range parameters, and *ϕ*_2_ > 0 and *λ*_2_ > 0 are the long range spatial and temporal range parameters, respectively. Further, *σ*^2^ > 0 is a spatio-temporal variance parameter, and 0 < *α* < 1. Hence, the full model for *z*(**s**, *t*) includes parameters 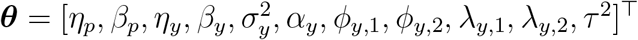

### 4.4 Disturbance

Predictions of forest carbon *z*(**s**, *t*), as well as model components *p*(**s**, *t*) and *y*(**s**, *t*), are readily available for pixels within well aggregates, and are obtained via their posterior predictive distributions. To understand baseline levels of forest area across the entire study domain, *p*(**s**, *t*) was estimated at each TCC pixel location in the first year (*t* = 1). For a given area such as a county, with geographic area denoted as 𝒮, the total forest area is determined as ∑_**s**∈𝒮_ *p*(**s**, 1), which is used as a baseline to quantify percent forest area disturbed for each county.

To assess instances of predicted forest disturbance at a location **s**, a threshold criteria is implemented based on estimated values of *µ*_*p*_(**s**, *t*). Specifically, for location **s** and time *t*, the probability of disturbance is calculated as *d*(**s**, *t*) = *µ*_*p*_(**s**, *t* − 1) *×* (1 − *µ*_*p*_(**s**, *t*)), which represents the probability that location **s** was forested at time *t* − 1 and not forested at time *t*. A location **s** is determined to have been disturbed if *d*(**s**, *t*) > 0.5 for some *t* > 1. The selection of this threshold seeks to balance identification of marginal disturbances associated with natural gas development while filtering minor disturbances associated with natural forest processes.

### 4.5 Opportunity Cost

To assess opportunity costs of disturbed well aggregate areas, an Autoregressive Moving Average (ARMA) model is fit using TCC values from undisturbed well areas to estimate carbon values for disturbed pixels. Specifically, TCC values from 10,000 undisturbed locations were used to fit the ARMA model, which was then used to predict undisturbed TCC values 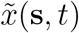 for disturbed locations **s** for all *t* > 1. These predicted TCC values 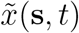 represent the undisturbed scenario for location **s**, which is used to calculate 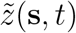 following models described in Section 4.3 using 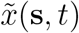 in place of *x*(**s**, *t*). Since the exact time of disturbance is not known, the opportunity cost for disturbed location **s** is taken as the difference in disturbed and undisturbed forest carbon in the final year, and is calculated as 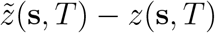.

## Acknowledgments

The findings and conclusions in this publication are those of the authors and should not be construed to represent any official USDA or U.S. Government determination or policy. The information is distributed solely for the purpose of pre-dissemination prior to peer review under applicable information quality guidelines. It has not been formally disseminated by the Forest Service. It does not represent and should not be construed to represent any agency determination or policy.

## References

Mary Beth Adams, W. Ford, Thomas Schuler, Melissa Thomas-Van Gundy, and Gundy. Effects of natural gas development on forest ecosystems. WV Research Wildlife Biologist, 01 2011.

William A. Bechtold and Paul L. Patterson, editors. The enhanced forest inventory and analysis program: national sampling design and estimation procedures. U.S. Department of Agriculture, Forest Service, Southern Research Station, Asheville, NC, 2005.

Kristin M. Carter, John A. Harper, Katherine W. Schmid, and Jaime Kostelnik. Unconventional natural gas resources in pennsylvania: The backstory of the modern marcellus shale play. Environmental Geosciences, 18(4):217–257, 12 2011. ISSN 1075-9565. doi: 10.1306/eg.09281111008. URL https://doi.org/10.1306/eg.09281111008.

Nicholas Coops, Liam Irwin, Harry Seely, and Spencer Hardy. Advances in laser scanning to assess carbon in forests: From ground-based to space-based sensors. Current Forestry Reports, 11, 01 2025. doi: 10.1007/s40725-024-00242-4.

Alice Favero, Adam Daigneault, and Brent Sohngen. Forests: Carbon sequestration, biomass energy, or both? Science Advances, 6(13):eaay6792, 2020. doi: 10.1126/sciadv.aay6792. URL https://www.science.org/doi/abs/10.1126/sciadv.aay6792.

Shawn Grushecky, Kevin Harris, Michael Strager, Jingxin Wang, and Anthony Mesa. Land cover change associated with unconventional oil and gas development in the appalachian region. Environmental Management, 70:869–880, 08 2022a. doi: 10.1007/s00267-022-01702-y.

Shawn T. Grushecky, F Christian Zinkhan, Michael P. Strager, and Timothy Carr. Energy production and well site disturbance from conventional and unconventional natural gas development in west virginia. Energy, Ecology and Environment, 7(4):358–368, apr 2022b. doi: 10.1007/s40974-022-00246-5. URL https://doi.org/10.1007/s40974-022-00246-5.

Kevin Harris. Disturbance related to unconventional oil and gas development in the appalachian basin. 2020. URL https://api.semanticscholar.org/CorpusID:220126478.

Ian Housman, Karen Schleeweis, Josh Heyer, Bonnie Ruefenacht, Stacie Bender, Kevin Megown, Wendy Goetz, and Seth Bogle. National land cover database tree canopy cover methods v2021.4. GTAC-10268-RPT1. Salt Lake City, UT: U.S. Department of Agriculture, Forest Service, Geospatial Technology and Applications Center, 2023.

Nathan F. Jones and Liba Pejchar. Comparing the ecological impacts of wind and oil & gas development: A landscape scale assessment. PLOS ONE, 8(11):1–12, 11 2013. doi: 10.1371/journal.pone.0081391. URL https://doi.org/10.1371/journal.pone.0081391.

Nathan F. Jones, Liba Pejchar, and Joseph M. Kiesecker. The energy footprint: How oil, natural gas, and wind energy affect land for biodiversity and the flow of ecosystem services. BioScience, 65(3):290–301, 01 2015. ISSN 0006-3568. doi: 10.1093/biosci/biu224. URL https://doi.org/10.1093/biosci/biu224.

Jessica Lovering, Marian Swain, Linus Blomqvist, Rebecca R. Hernandez, and Lalit Chandra Saikia. Land-use intensity of electricity production and tomorrow’s energy landscape. PLoS ONE, 17(7), 7 2022. doi: 10.1371/journal.pone.0270155.

Paul B. May and Andrew O. Finley. Spatial-temporal models for forest inventory data, 2025. URL https://arxiv.org/abs/2503.16691.

Robert I. McDonald, Joseph Fargione, Joe Kiesecker, William M. Miller, and Jimmie Powell. Energy sprawl or energy efficiency: Climate policy impacts on natural habitat for the united states of america. PLOS ONE, 4(8):1–11, 08 2009. doi: 10.1371/journal.pone.0006802. URL https://doi.org/10.1371/journal.pone.0006802.

Elliot S Shannon, Andrew O Finley, Grant M Domke, Paul B May, Hans-Erik Andersen, George C Gaines III, and Sudipto Banerjee. Toward spatio-temporal models to support national-scale forest carbon monitoring and reporting. Environmental Research Letters, 20(1):014052, dec 2024. doi: 10.1088/1748-9326/ad9e07. URL https://doi.org/10.1088/1748-9326/ad9e07.

Elliot S. Shannon, Andrew O. Finley, Paul B. May, Grant M. Domke, Hans-Erik Andersen, George C. Gaines III, Arne Nothdurft, and Sudipto Banerjee. Leveraging national forest inventory data to estimate forest carbon density status and trends for small areas, 2025. URL https://arxiv.org/abs/2503.08653.

E.T. Slonecker, L.E. Milheim, C.M. Roig-Silva, A.R. Malizia, D.A. Marr, and G.B. Fisher. Landscape consequences of natural gas extraction in bradford and washington counties, pennsylvania, 2004–2010. URL http://pubs.usgs.gov/of/2012/1154.

Anne M. Trainor, Robert I. McDonald, and Joseph Fargione. Energy sprawl is the largest driver of land use change in united states. PLOS ONE, 11(9):1–16, 09 2016. doi: 10.1371/journal.pone.0162269. URL https://doi.org/10.1371/journal.pone.0162269.

U.S. Energy Information Administration. Appalachian basin drives growth in u.s. natural gas production, 2021. URL https://www.eia.gov/todayinenergy/detail.php?id=48976. Accessed June 9, 2025.

J A Westfall, J W Coulston, G G Moisen, and H E Andersen, editors. Sampling and estimation documentation for the Enhanced Forest Inventory and Analysis Program: 2022. General Technical Report NRS-207, U.S. Department of Agriculture, Forest Service, Northern Research Station, Madison, WI, 2022.

John Young, Kelly O. Maloney, E. Terrence Slonecker, Lesley E. Milheim, and David Siripoonsup. Canopy volume removal from oil and gas development activity in the upper susquehanna river basin in pennsylvania and new york (usa): An assessment using lidar data. Journal of Environmental Management, 222:66–75, 2018. ISSN 0301-4797. doi: 10.1016/j.jenvman.2018.05.041. URL https://www.sciencedirect.com/science/article/pii/S0301479718305668.

